# Mechanisms of ecological divergence with gene flow in a reef-building coral on an isolated atoll in Western Australia

**DOI:** 10.1101/2021.08.08.455318

**Authors:** L Thomas, JN Underwood, NH Rose, ZL Fuller, ZT Richards, L Dugal, C Grimaldi, IR Cooke, SR Palumbi, JP Gilmour

## Abstract

Understanding the mechanisms driving phenotypic variation in traits facing intensified selection from climate change is a crucial step in developing effective conservation and restoration initiatives. This is particularly true for reef-building corals, which are among the most vulnerable to climate change and are in dramatic decline globally. At the Rowley Shoals in Western Australia, the prominent reef flat becomes exposed on low tide and the stagnant water in the shallow atoll lagoons heats up, creating a natural laboratory for characterising the mechanisms that control phenotypic responses to different environments. We combined whole genome re-sequencing, common garden heat stress experiments, transcriptome-wide gene expression analyses, and symbiont metabarcoding to explore the mechanisms that facilitate survival in contrasting habitat conditions. Our data show that, despite high gene flow between habitats, spatially varying selection drives subtle shifts in allele frequencies at hundreds of loci. These changes were concentrated into several islands of divergence spanning hundreds of SNPs that showed strong linkage disequilibrium and were associated with a coordinated increase in minor allele frequencies in corals taken from the lagoon habitat, where the range of environmental conditions is greatest. Common garden heat stress assays showed individuals from the lagoon exhibited higher bleaching resistance than colonies from the reef slope, and RNAseq identified pronounced physiological differences between the corals from the two habitats, primarily associated with molecular pathways including cell signalling, ion transport and metabolism. Despite the pronounced physioloigical and environmental differences between habitats, metabarcoding of the *Symbiodiniaceae* ITS2 region revealed all colonies to be associated exclusively with the genus *Cladocopium*, with no detectable differences between habitats. This study contributes to the growing number of studies documenting the complex mechanisms that facilitate coral survival in extreme environments, and showcases the utility of combining multiple sequencing techniques to unravel complex climate-related traits.

## Introduction

The impacts of anthropogenic climate change and associated extreme weather events are devastating natural ecosystems around the world^1–3^. Coral reefs are especially threatened, and it is uncertain whether reef-building corals can keep pace with this unprecedented rate of environmental change^4,5^. Fortunately, most coral species have large effective population sizes with broad geographic ranges that span strong environmental gradients over which their phenotypes vary^6,7^. As a result, they tend to harbour an abundance of phenotypic and genetic variation in fitness-related traits that are facing intensified selection with climate change, such as thermal tolerance^8–15^. The resulting ecological divergence among populations and associated levels of connectivity will underlie the capacity of species to adapt to rapid environmental change over the coming decades^16–18^.

Thermal tolerance in corals is a highly heritable and polygenic trait^19–23^, governed by a complex interplay between host genetics and symbiont community composition^9^. Studies documenting transcriptome-wide changes in gene expression regularly report hundreds of genes that respond to environmental stress and that are associated with a diverse range of cellular pathways^14,21,24–26^. Consistent with these patterns, tens to hundreds of loci are often reported in *F*_ST_ based outlier analyses among corals across large environmental gradients^23,27,28^. However, when gene flow is high, as in most broadcast-spawning corals, adaptation favours the tight linkage of small effect loci into regions of reduced genetic distance and recombination^29,30^. In this case, specific genomic regions may play a particularly important role in driving evolutionary change^22,31^. Identifying genomic regions that control complex climate-related traits is central to developing a mechanistic and predictive understanding of climate-change resilience in corals. In turn, this information can be used to help design effective spatial management strategies aimed at protecting areas of reef that harbour high-levels of stress tolerant alleles, as well as improve more proactive management interventions such as reef restoration and assisted gene flow initiatives^32,33^.

Variation in thermal tolerance occurs across large spatial scales (100s of kilometres)^31,34–36^, where populations are exposed to contrasting environmental conditions, but also on extraordinarily fine-spatial scales (100s of metres) across different habitats on the same reef^8,9,14,37–40^. A combination of complex bathymetry, hydrodynamics and water quality can expose some coral communities to more variable or extreme environmental conditions than other areas of reef ^38^. These relatively extreme environments select for stress resistance and can offer important insight into the mechanisms of climate change resilience in corals^9,10,19,23,38^. These populations from variable or more extreme habitats are a natural asset for reef management, yet we have a limited understanding of the mechanisms at play, which impedes our ability to take proactive steps towards effectively managing coral reefs for resilience against climate change^32,33^.

The lagoon at each of the three reef atolls at Western Australia’s Rowley Shoals is a highly variable and at times extreme habitat that supports extensive coral growth. These atolls have a prominent reef flat that greatly restricts any flushing or exchange of water with the open ocean. As a result, coral populations in the lagoon are regularly exposed to different temperature and flow dynamics than those on the reef slope (Figure 1, Table S1, ESM2). Despite these contrasting environmental conditions, some coral species, such as the wide-spread and ecologically important *Acropora tenuis*, are common in both habitats. Here, we combine whole genome re-sequencing, common garden heat stress experiments, transcriptome-wide gene expression analyses, and symbiont metabarcoding to explore the mechanisms of fine-scale ecological divergence in a dominant reef-building coral on a remote atoll in northwest Australia.

**Figure 1.**
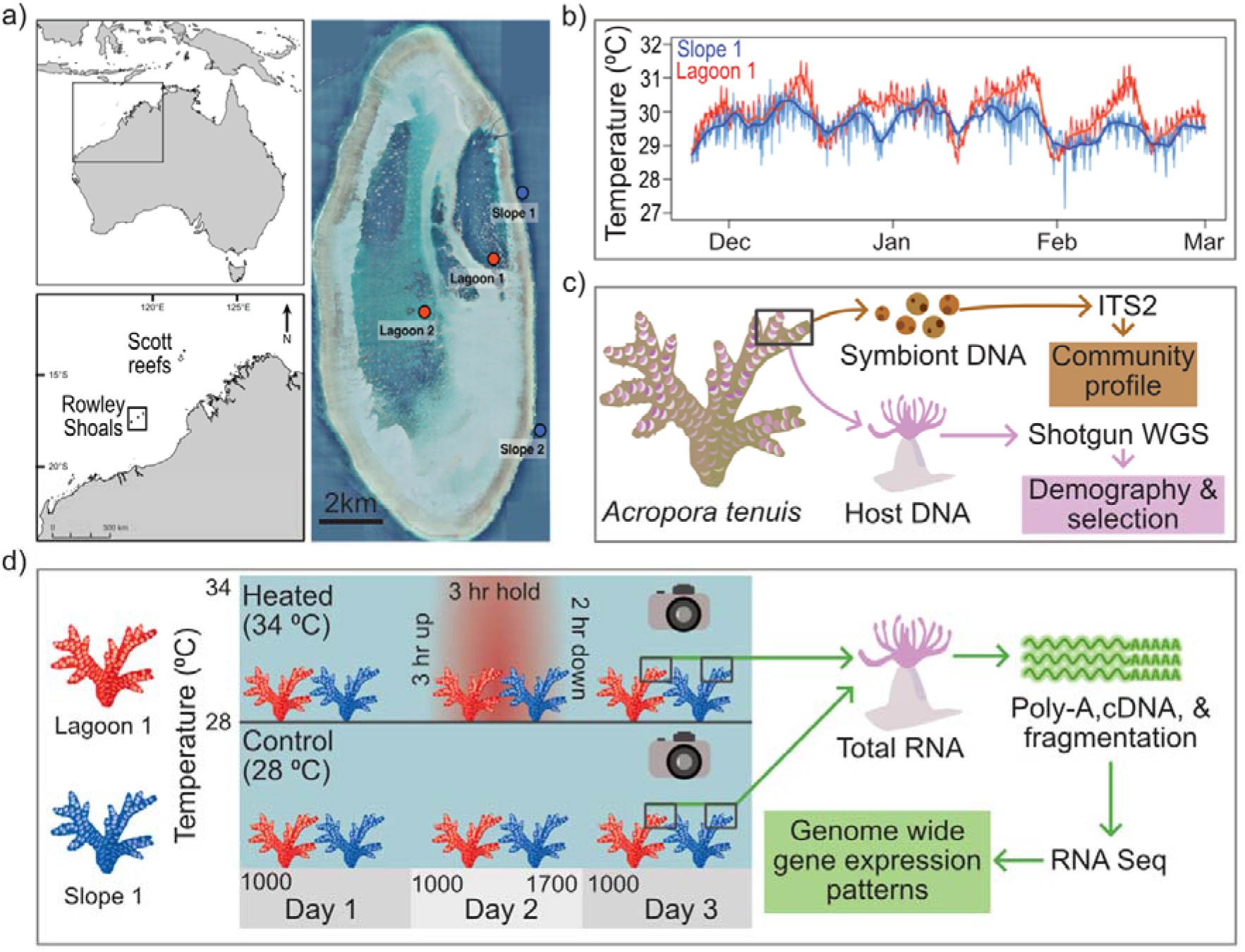
Sample sites, experimental design, and sequencing methods. **A)** Map of sampling locations in lagoon and slope habitats at Clerke Reef in the Rowley Shoals; **B)** Time-series temperature plot of temperaure in lagoon and slope sites across 2017/18 summer; **C)** Tissue samples for whole genome sequencing (WGS) and ITS2 metabarcoding were collected from Acropora tenuis at two lagoon (n=20) and two slope (n=20) sites (n=80 total). Samples were also collected from the reef slope habitat at Scott Reef (n=10) for WGS to place levels of divergence among habitats in a broader biogeographic and evolutionary context; **D)** Experimental set up for common garden acute heat stress assays. Colony fragments used in experimental heat stress and RNA-seqeuncing (n=10 per habitat) were collected from Slope 1 and Lagoon 1.

## Materials and Methods

### 1. Demographic history, population structure, and signatures of selection with whole genome re-sequencing

#### Sample collection

*Acropora tenuis* forms a species complex in Western Australia with morphologically cryptic colonies that occur in sympatry but spawn in either spring or autumn^41^. To avoid confounding our data by including two distinct genetic lineages, we screened 225 colonies collected across 4 sample sites (2 lagoon and 2 slope) at Clerke Reef with a panel of nine microsatellite loci that can differentiate between the autumn and spring spawning cohort^41^. Samples were collected along a 200m transect from colonies separated by at least 1.5m and preserved in ethanol immediately after collection. DNA was extracted from samples using Qiagen DNA Blood and Tissue kit. Allelic variation was measured at nine microsatellite loci^41^ and Bayesian admixture analysis in *structure*^42^ was used to assign individuals to either spring or autumn cohort using known reference colony genotypes (Table S2). From this analysis we identified 20 individuals from the spring cohort from two lagoon and two slope sites at Clerke Reef (Figure 1A) that we used for downstream sequencing and analyses. The spring cohort was chosen because it was common in both habitats, in contrast to the autumn spawner which was more dominant on the reef slope. We also included 10 samples from the reef slope of the neighbouring Scott reef system to place differences among lagoon and slope corals at Clerke Reef into a broader geographic and evolutionary context.

#### Genotype likelihoods

A recently developed reference assembly for *A. tenuis* is one of the most complete genome assemblies available for corals to date^43^. However, it remains fragmented and lacks chromosome-scale resolution. In an attempt to improve this assembly, we used *ragoo*^44^ to generate *pseudo* chromosomes for the *A. tenuis* assembly (*aten_final_0.11.fasta*) using the recently published chromosome-scale *Acropora millepora* genome assembly^22^. *Ragoo* is a reference-guided scaffolding method that orders and orients assembly scaffolds and contigs to a single reference genome^44^. We ran *ragoo* with default parameters using *minimap2* as the aligner^45^. A total of 407Mb of sequence were localized, corresponding to 83.6% of *A. tenuis* contigs, 94% of which fell within the 14 *A. millepora* chromosomes. Gene coordinates from the reference *A. tenuis* assembly were lifted over to the pseudo-chromosomes using the *ragoo* utility script *lift_over.py* included in the software.

Shotgun libraries were prepared and individually barcoded using Nextera Flex Library Preparation Kits (Illumina) and pooled together for sequencing on a NovaSeq 6000 (S1, 300 cycles). Of the 90 samples we processed, five samples from Rowley Shoals did not produce libraries of sufficient quality for sequencing and were excluded from this study. Raw FASTQ files were processed using *fastp*^46^ for adapter trimming, quality filtering (>=Q15), and down sampling samples to a maximum of 10M paired-end reads. This was done to ensure equal sequencing depth across samples. After initial QC, reads were mapped with *bwa mem* to the *A. tenuis* pseudo-chromosome-scale assembly (*aten.chr.fasta*). We used *samtools*^47^ to sort and index and *picardtools* (http://broadinstitute.github.io/picard) to mark and remove duplicate reads. *Samtools* was also used to calculate sequencing depth of each sample at each position across the genome. We used *angsd*^48^ to estimate the site frequency spectrum (SFS) for each population based on genotype likelihoods. We masked sites minQ < 20, minMapQ < 30, sites with less than 1/3 and greater than 2x the mean read depth for each sample site. We also filtered out loci with missing data in more than half of the samples per site.

#### Demographic history

To provide an overall demographic and evolutionary context of the Rowley Shoals, we calculated genome-wide estimates of Tajima’s D using *realSFS*^48^. A positive Tajima’s D indicates an absence of low frequency alleles, suggestive of a recent bottleneck. These analyses were based on the unfiltered folded site frequency spectrum (ancestral allele state unknown) using 1-kb non-overlapping windows. We also sequenced one individual from the Rowley Shoals reef slope to ~46x coverage and applied the partially sequentially Markovian coalescent (PSMC’) method implemented in MSMC^43,49^. *Snpable* (http://lh3lh3.users.sourceforge.net/snpable.shtml) was first used to mask all regions of the genome where it was not possible to obtain unambiguous read mappings. Variant calling was performed for deep sequenced samples using *samtools* and *bcftools*. A distribution of MSMC estimates was obtained by generating 100 bootstrap runs, each of which involved recombining a random subsampling of 500kb sized chunks to produce 20 large scaffolds of length 10Mb. Raw outputs from MSMC were converted to real values using a mutation rate of 1.86e^−8^ events per base per generation and a generation time of 5 years. This choice of generation time is consistent with the fast growth rate and relatively high turn-over of *Acropora*^43,50^ and the value for μ was estimated based a divergence time of 15.5Mya between *A. tenuis* and *A. digitifera*^46^ using synonymous substation rates for one to one orthologs as described in Cooke et al^39^. The values of *N_e_* and time we infer depend on our assumptions of μ and the generation time, and we note that any uncertainty in these parameters is reflected in the absolute values, but will not affect the qualitative patterns of population size changes.

#### Population structure and signatures of selection

To explore patterns of genetic differentiation between habitats, we excluded variant sites with maf < 0.05 and snp_value > 1e-6 and used *pcangsd*^51^ to output a covariance matrix that was used as an input to the *R* function *eigen* to carry out PCA. *Pcangsd* also estimates the optimal number of clusters and uses a matrix factorization algorithm to estimate admixture proportions for each sample. To estimate divergence across the mitochondrial genome, we also mapped reads to the complete *A. tenuis* mitochondrion^52^ (18,659 base pairs) and extracted consensus sequences for each sample with *samtools*. We aligned these consensus sequences using a Muscle alignment in *Geneious* and then built a TCS network in *Popart*^53^.

When gene flow is high, as in most broadcast-spawning corals, adaptation favors the tight linkage of small effect loci into regions of reduced genetic distance and recombination^29,30^. In such cases, even for complex polygenic traits, specific genomic regions may play a particularly important role in driving evolutionary change^22,31^. To this end, we searched for signatures of selection in coral from the warm lagoon habitat at Clerke Reef by calculating *F*_ST_ between habitats using 100-kb windows with 10-kb steps and identified empirical outliers as those occurring in the top 0.01% of windows genome-wide. This sliding window approach allowed us to hone in on large genomic regions of elevated differentiation, or islands of divergence, rather than focusing on any single variant alone. These analyses were done using the unfiltered SFS for each habitat. We tested for strong linkage disequilibrium across outlier regions using *ngsLD*^54^, which estimates pairwise linkage among variant sites based on genotype likelihoods. Finally, we used a population-blind approach to identify targets of selection using the *-selection* flag in *pcangsd*. These analyses were run with the Scott Reef samples removed from the dataset. The program looks for SNPs with a distribution that exceeds expectations under neutrality along the first principal component. We calculated a selection statistic for each SNP (MAF > 0.05) along each chromosome separately, and calculated outlier probabilities using the *pchisq* function (two-tailed mode) in R.

### 2. Physiological differences with acute experimental heat stress and RNA-seq

#### Common garden heat stress experiment

To explore the physiological differences between coral from the two habitats, we combined common garden acute heat stress experiments with RNA sequencing. Replicate fragments (n=2) from 15 colonies of *A. tenuis* from the lagoon (L1) and slope (S1) habitats at Clerke Reef were exposed to ambient (28°C) and heated (34°C) treatments using a portable temperature-controlled flow-through seawater system aboard our research vessel RV Solander. After 24hrs of acclimation in the tanks, nubbins were exposed to acute heat stress, which consisted of a 3-hour temperature ramp to 34°C starting at 10:00 am, followed by a three-hour hold at 34°C and then a two-hour ramp back down to ambient temperatures^8,9,55^. The next morning, twenty hours after the acute heat stress assay began, replicate nubbins from control and heated treatments were photographed to quantify pigment loss using the photographic method^56,57^, which uses shifts in intensity along the Red spectrum as a proxy for chlorophyll loss. Samples were then flash frozen in liquid nitrogen for gene expression analyses. Although an acclimation phase of one day in a common garden was designed to remove gene expression changes due to fragmentation or acute environmental responses, it was likely not long enough to erode any long-term plastic effects on gene expression.

#### Genome-wide gene expression patterns

Total RNA was extracted from 40 samples (10 heated and 10 control from each habitat) using a QIAGEN RNAeasy mini kit and stranded poly-A RNA libraries were prepared using Agilent’s SureSelect library preparation kit. The protocol includes a polyA enrichment followed by fragmentation, reverse transcription, second strand synthesis with UTP, adapted ligation and 12 cycles of UDG for indexing. Library QC was performed using TapeStation 4100 and were re-pooled based on iSeq data before being sequenced on a NovaSeq SP flow cell in 2 x 50 cycles format to yield approximately 10M read-pairs per sample at Genomics WA. Raw FASTQ files were processed using *fastp*^46^ for adapter trimming and quality filtering (>=Q30) and reads were then mapped using *hisat*^58^ to the *A. tenuis* genome. As a first quality control step and to avoid including multiple spawning lineages in our analyses of gene expression, we used *samtools* to call variant sites and *prcomp* in R to carry out principal component analysis. Once we isolated a single genetic lineage (spring), differential gene expression analyses were carried out in *DESeq2*^59^ using a raw counts matrix. Differentially expressed genes were identified using Wald tests after adjusting for multiple comparisons (*p.adj* < 0.05). Functional enrichment based on Gene Ontology (GO) terms was assessed using a two-tailed Mann-Whitney U test in *gomwu*^60^. To further quantify the magnitude of response to heat stress, we used Discriminant Analysis of Principle Components and measured the magnitude of shift along the first principle component as a quantitative measure of the overall gene expression response^39^.

### 3. Symbiont community profiling with ITS2 metabarcoding

Complex interactions between the coral host and the endosymbiotic algae (Family *Symbiodiniaceae*^61^) are superimposed on the coral genetics that drive variation in environmental tolerance^62–64^. In Western Australia, *Acropora* have a high level of affinity to the genus *Cladocopium*^65^, so to determine if corals from different habitats at the Clerke Reef associate with different *Cladocopium* types we applied metabarcoding of the ITS2 region of the ribosomal DNA for a subset of the colonies (n= 65) collected across the four sample sites at Clerke Reef. The internal transcribed spacer 2 (ITS2) region was amplified using the ITSD (5⍰-GTG AAT TGC AGA ACT CCG TG-⍰3) and ITS2-rev2 (5⍰-CCT CCG CTT ACT TAT ATG CTT-⍰3 primer pair. PCR was carried out in 30 μl reactions containing 15μl 2x Phusion Master Mix, 0.5μl of each primer (10μM) modified with Nextera Illumina overhang sequences, 1.5μl of template DNA and water to volume. Thermal cycling conditions were as follows: denaturation at 98°C for 30 s, followed by 35 cycles of 10 s at 98°C, 30 s at 60°C and 30 s at 72°C and a final extension at 72°C for 5 min (performed on a Mastercycler Nexus Gradient (Thermofisher Scientific). Amplicons underwent a second round of PCR to attach flow cell adapters and unique barcodes, and were then pooled at equal molar concentrations for sequencing on an Illumina MiSeq using a V2 600 cycle paired end kit at the Australian Genome Research Facility (AGRF). De-multiplexed reads were processed through *symportal*^66^, which was designed specifically for the analysis of *Symbiodiniaceae* ITS2 metabarcoding data.

## Results and Discussion

### 1. Genotype likelihoods and demographic history

We used a low coverage whole genome sequencing approach and mapped 573,937,253 paired-end (250 cycles) sequence reads from 85 colonies of *A. tenuis* (spring spawning lineage^41^) to our *A. tenuis* pseudo-chromosome assembly, achieving a mean genome sequencing coverage of approximately 4.2x (+/− 0.10 SE) per sample (Figure S2; Table S3). We also mapped reads to the complete *A. tenuis* mitochondrion and achieved a mean coverage of 313x (+\− 18.27 SE) per sample. Genotype likelihoods were calculated across 282,878,358 sites that met our filtering criteria for coverage and quality. Initial screening of samples identified three outlier colonies (two of which appeared to be clones) that were collected from one of the lagoon sites (L2) and that were removed from downstream analyses (Figure S3). These samples likely represent either mis-IDs or cryptic species that were not resolved with the panel of microsatellite markers. Estimates of Tajma’s D at the Rowley Shoals showed a strong positive shift relative to Scott Reef, indicative of a recent bottleneck (Figure 2A). Demographic changes from 50kya to 5Mya inferred from MSMC based on a single high-coverage individual showed a steady decline in *N_e_* at the Rolwey Shoals following a broad peak between ~100k and 1M years ago (Figure 2B). This peak in effective population size ~150,000 years before present is similar to other estimates for this species from the Indo-Pacific^43,50^.

**Figure 2.**
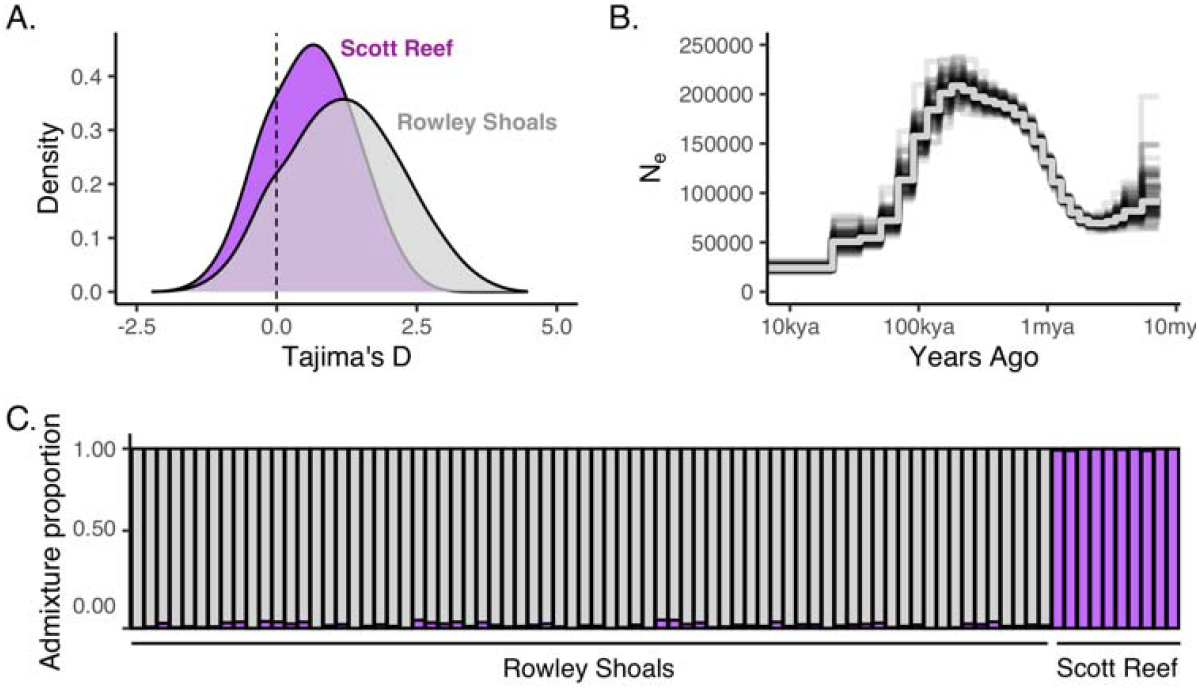
Demographic history and regional population structure in A. tenuis from the Rowley Shoals and the Scott reef system. **A)** Density plot of Tajima’s D across the genome for Rowley Shoals (grey) and Scott Reef (purple); **B)** Changes in population size through time the Rowley Shoals using PSMC’ from deep sequencing (45x) of a single colony. Individual black lines show bootstrap replicates and solid grey line shows the average; and **C)** Admiture proportions of samples from Scott Reef (purple) and Rowley Shoals (grey) based on genotype likelihoods using ~6.5M variant sites.

### 2. Population differentiation and signatures of selection

Multiple clustering methods based on the nuclear data indicated strong genetic subdivision between Scott Reef and Rowley Shoals (Global *F*_ST_ = 0.25; Figure 2C, Figure S3). We did not, however, observe any clear geographic clustering of samples based on the complete mitochondrial genomes (Figure S4). Considering the strong divergence across the nuclear genome, a lack of clear structure across the mitochondrion likely reflects historical introgression and/or the low mutation rates characteristic of cnidarians^67^. Within Clerke Reef, we detected low levels of nuclear divergence among samples (Weighted *F*_ST_ = 0.007), with no spatial clustering of colonies by habitat (Figure 3A). Principle component analysis based on 5,493,423 SNPs (maf > 0.05) showed all samples to be admixed across habitats, indicative of high gene flow (Figure 3A). This lack of differentiation between habitats was not surprising considering the spatial proximity of sample sites and the broadcast spawning life history strategy of *A. tenuis*. It is also consistent with patterns of gene flow between habitats in the congener *Acropora digitifera* from the Rowley Shoals^68^.

**Figure 3.**
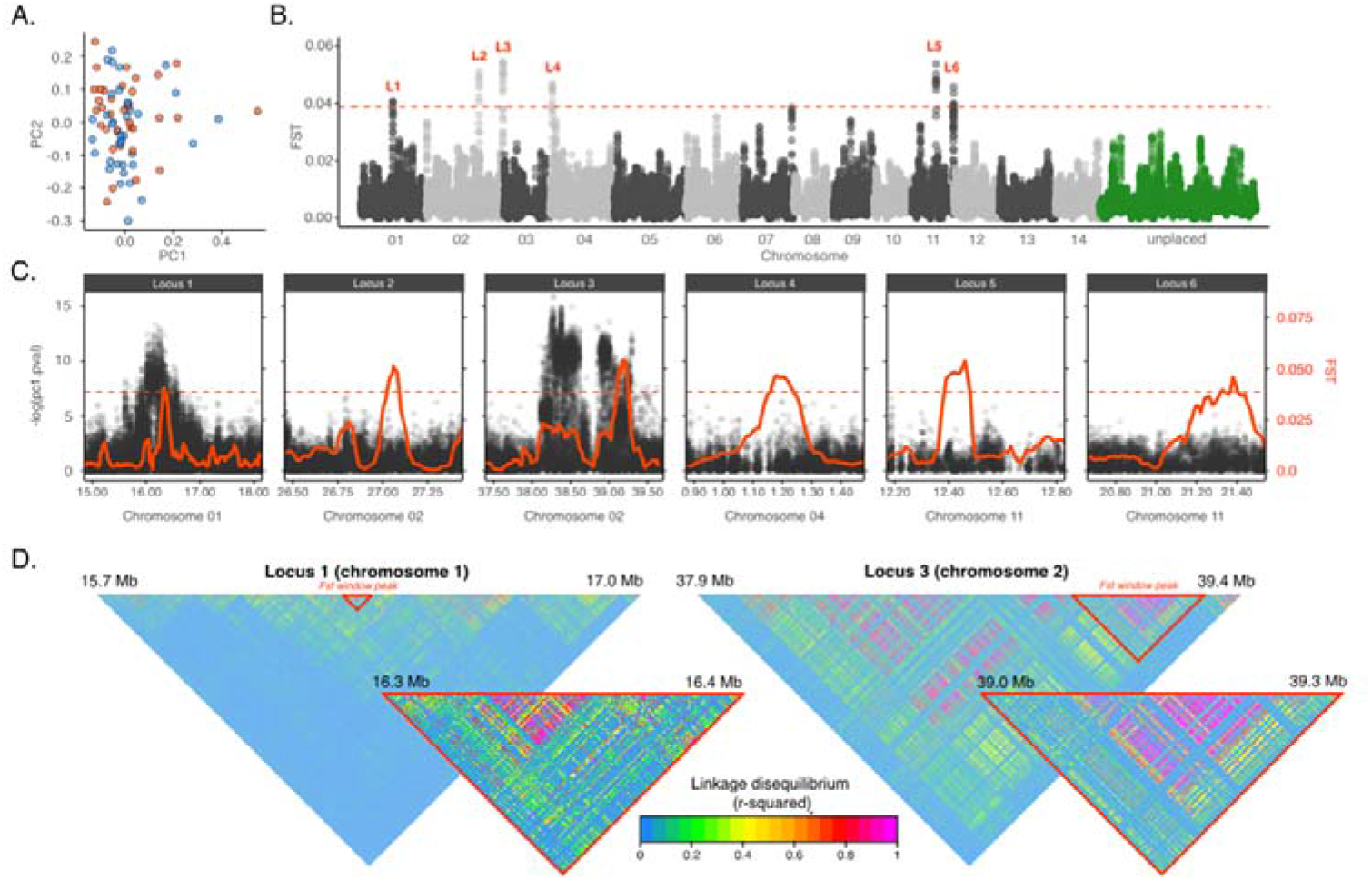
Genome-wide patterns of differentiation in A. tenuis from lagoon and slope habitats. **A)** Scatter plot of the first two principal components for colonies from lagoon (orange) and slope (blue) habitats based on genotype likelihoods across 5,493,423 SNPs (maf > 0.05); **B)** Manhattan plot of the patterns of F_ST_ across the genome using 100kb windows (10kb step) based on the unfiltered SFS across 433,290,581 sites. Alternating colours indicate different chromosomes. Each point is the average F_ST_ of all SNPs in that window. Top 0.01% windows (red dotted line) highlighted in red; **C)** Zoomed in plots of F_ST_ (red line-100kb windows), and the population-independent selection coefficients for each SNP (black points) surrounding outlier regions; **D)** Heatmap of linkage disequilibrium (r2) across Locus 1 and Locus 3. Warmer colors indicate higher linkage among SNPs. Notice the two window peaks overlap regions of high LD.

A lack of any clear genetic structure between habitats at Clerke Reef suggests that local recruitment comes from a mixed larval pool. In such cases, environmental heterogeneity gives rise to patterns of spatially varying selection where different habitats select for genotypes from a common gene pool that match the local conditions^69^. In turn, this results in an overall increase in adaptive genetic variation in the broader metapopulation^70^. Alleles maintained at moderate frequencies through balancing selection are particularly important for adaptation to divergent habitats^22,71,72^. In contrast to directional selection, which reduces genetic variation by fixing beneficial alleles, balancing selection maintains genetic variation within a species and plays a crucial role in adaptation in species with high gene flow and that span strong environmental gradients^22,73,74^.

Although a lack of population structure between coral from the lagoon and slope suggests regular dispersal between habitats (Figure 3A), we identified several regions of the genome that showed high levels of differentiation (*F*_ST_) and were strong candidates for selection (Figure 3B). The most extreme values genome-wide overlapped 6 islands of divergence spread across 4 different chromosomes (Figure 3C). Mean *F*_ST_ across these windows was 0.047, compared to a genome-wide average across all windows of 0.007. Our population-blind approach using PCAngsd identified two of these regions (Locus 1 and Locus 3) to include a large number of SNPs showing signs of selection with distributions that exceeded what was expected under neutrality along the first principal component (Figure 3C).

Estimates of linkage disequilirium (LD) across Locus 1 and Locus 3 showed *F*_ST_ window peaks to span strong linkage blocks and associated with coordinated changes in allele frequencies across hundreds of loci (Figure 3D). Across these regions, we observed a significant increase in minor allele frequencies (Welch Two Sample t-test, p<0.001) in lagoon colonies relative to those from the reef slope (Figure S5). Allele frequencies seemed to be concentrated at a particular value in each habitat, a further sign of reduced recombination and supporting the observed patterns of LD. Interestingly, when we carried out a PCA across Locus 1 and Locus 3, samples cluster strongly into three distinct groups along the first principle component, likely representing the three possible genotypes for each haplotype block (Figure S6). Notably, the two dominant genotypes showed different proportions in lagoon and slope samples.

Locus 1 directly overlapped genic regions with homologues to two Tolloid-like proteins involved in zinc ion binding and development, but also included a number of genes within the broader linkage group that had homology to classic stress response proteins, including E3 ubiquitin protein ligases (Lin41/Cop1), Tumor necrosis factor receptors (TFIP8, TRAF2, TRAF3), and heat shock proteins (HSP70) and chaperones (DnaJ homologs) (Table S3), but were not enriched for any specific molecular function (p.adj < 0.05). Locus 3 *F*_ST_ window peak directly overlapped genic regions with homology to Phosphatidylinositol-glycan biosynthesis class A proteins. Several other genes fell within the broader linkage block upstream of Locus 3, including a Death-associated protein kinase 3 (DAP kinase 3), a serine/threonine kinase (Kalirin), Myosin light chain kinase (MLCK), a heat shock protein (HSP75) and a NACHT domain-containing protein (NLRP5) (Table S4). Genes in this broader linkage group upstream of Locus 3 were highly enriched for ATP binding (GO:0005524, p.adj=0.00026), suggesting a role in enzyme regulation and environmental response pathways.

Areas of high linkage and reduced recombination are often associated with chromosomal rearrangements, such as inversions, which play a key role in driving phenotypic variation and local adaptation^30,75^. One of the classic examples of this comes from mimetic wing patterning in *Helioconus* butterfly, where several inversions along a 200 kb region of the genome under strong linkage disequilibrium are associated with different wing patterning^30^. Whether the patterns observed here are actual inversions remain unclear, and future research that utilizes long read sequencing (e.g., Oxford nanopore or PacBio technology) will help elucidate the role of structural rearrangements in reef coral adaptation to climate change.

Gene flow in *A. tenuis* between habitats is high across most of the genome, but barriers to gene flow exist. A number of genomic regions showed strong habitat-specific shifts in allele frequencies across hundreds of linked SNPs. This pattern is consistent with adaptation amidst gene flow, where beneficial genetic variants are consolidated into regions of reduced genetic distance and recombination^29,76^. Our data show that no single locus confers resilience to the warm lagoon, but in fact large coordinated changes in allele frequencies across large linkage blocks contribute to survival. It is important to note that if those allele frequency shifts confer resilience to certain environment stressors, then that shift must come at a cost, otherwise the alternate allele would reach fixation in the broader meta-population and there would be no variation in this trait. A better understanding of the fitness-related trade-offs of survival in marginal environments is essential future research, but points to growth, calcification and reproduction as primary costs of survival in marginal environments^10,15,77^.

### 3. Physiological differences between habitats

Coral from the reef slope were far more pigmented than lagoon corals at the onset of the experiment, but suffered greater pigment loss following short-term acute heat stress (Figure 4A). These patterns were consistent with our visual scoring method with the CoralWatch^®^ Coral Health Chart (Figure S7). Although we did not measure cell density or chlorophyll directly, our photographic method has been shown to correlate strongly with chlorophyll levels^57^. Reduced symbiont cell densities and chlorophyll levels have been directly linked to bleaching susceptibility in coral^78^, and maintaining lower levels appears to be one mechanism through which lagoon corals minimize photo-oxidative damage of a shallow high temperature environment.

**Figure 4.**
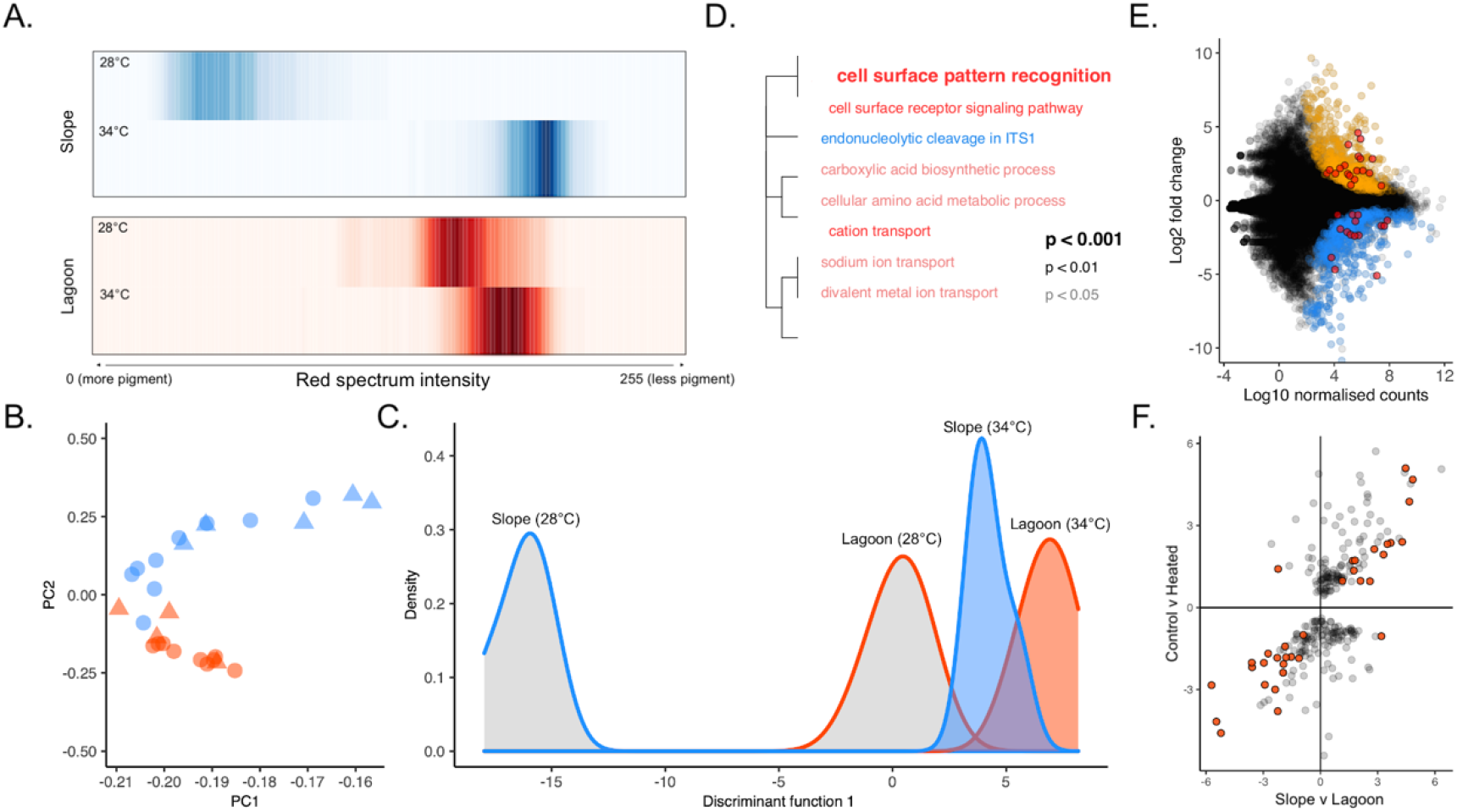
Physiological differences in A. tenuis from lagoon and slope habitats. **A)** Changes in pigmentation (Red channel intensity) of coral nubbins from lagoon (red) and slope (blue) habitats exposed to control (28°C) and heated (34°C) conditions. The darker colours indicate higher density of that pigment value. Values range from 0 (black) to 255 (white), and a positive shift to the right indicates a decline in pigment; **B)** Scatter plot of the first two principle components of normalized gene expression in A. tenuis in control (blue) and heated (red) treatments across 21,230 genes with a minimum mean depth per sample of 5. Circles denote samples from lagoon and triangles from reef slope; **C)** Density distribution of colonies along the first principle component and shows lagoon control samples to cluster closer with the heated treatments than with the other slope controls; **D)** Functional enrichment anlayses indicating enriched GO annotations for genes that showed baseline differences in expression between habitats; Blue is upregulated and red downregulated in lagoon samples; **E)** Regression of differentially expressed genes between habitats after two days acclimation at ambient temps. Each point represents a gene that showed significant differences in baseline expression between habitats. Red points were genes that form part of the heat stress response and were differnetially expressed between control and heated treatments across all samples; **F)** Linear regression of levels of log fold change in heat responsive genes between habitats, illustraing that patterns of gene expression under acute heat stress mirror expected changes in expression between habitats.

RNA-sequencing of samples from the common garden heat stress experiments revealed pronounced physiological differences between corals from the two habitats. In total, we mapped 462,429,066 paired end reads from 40 cDNA to the *A. tenuis* genome assembly. Initial screening of samples based on a panel 60,941 variant sites across the transcriptome revealed that one individual from the lagoon and five from the reef slope from our heat stress experiment belonged to the divergent autumn spawning lineage, and so we removed these samples from our analyses to ensure we focussed our analyses on a single gene pool (Figure S8). After these samples were removed, the RNAseq dataset confirmed the negligible divergence (mean *F*_ST_= −0.004) between habitats that was observed with the low-coverage WGS dataset (Figure S8).

Normalised gene expression profiles across 21,230 genes (Table S6) showed that lagoon corals held at ambient conditions clustered closely with samples from the heated treatments, and experienced a smaller overall transcriptomic shift in response to acute heat stress than corals from the slope (Figure 4). We identified 274 genes that differed in baseline levels of expression (p.adj<0.05) between colonies from the two habitats after short-term common garden acclimation (Figure 4C; Table S7). This gene list was enriched for Gene Ontology terms related predominantly to transport, metabolism and signalling (Figure 4D). Data from our acute heat stress assay showed approximately 12% (n=34) of the genes that differed in baseline expression between habitats were heat-responsive genes that were differentially expressed between heated and control treatments across all samples (Figure 4E, Table S8). This gene set followed intuitive expression profiles based on habitat; genes that were upregulated under acute heat stress had higher baseline expression in lagoon corals, and genes that were downregulated in response to acute heat stress had lower baseline expression in lagoon corals (Figure 4F). This gene set included classic stress response genes such as ubiquitin and zinc finger proteins and serine/threonine protein kinases (Table S9).

Corals are capable of pronounced physiological adjustment in response to heat stress, and individuals thriving in environments that are regularly exposed to environmental extremes often show pronounced differences in baseline levels of gene expression than adjacent colonies from more benign environments^19,39,79,80^. Higher baseline expression of stress response genes can promote stress tolerance and maintenance of cell homogeneity, while reduced expression can act to limit the negative physiological responses to stress^79^. Throughout the daily tidal cycle, corals in the lagoon at Clerke Reef experience warmer water and less current flow than their reef slope counterparts. Our data show that survival in such an environment requires a pronounced physiological shift across hundreds of genes. Through this physiological shift, lagoon corals are primed for stressors associated with the daily tidal cycle.

Levels of congruence were relatively low between differentially expressed genes from RNAseq dataset (habitats or heat stress) and the genomic islands of differentiation identified using our low-coverage WGS approach; few heat- or habitat-responsive genes occurred in regions of high genetic differentiation in the WGS datasets. The only exception to this was Locus 1 (Figure S9), with an *F*_ST_ window peak that directly overlapped *till1*, a Tolloid-like proteinase involved in zinc-finger binding and that showed strong changes in regulation under experimental acute heat stress (lfc: 2.62, p.adj: 5.8e-14) but that was not differentially expressed between habitats under control or heated conditions (lfc=0.46, p.adj: 0.78). It was also noteworthy that a number of other heat (n=7) or habitat (n=3) responsive genes occurred up- or down-stream of the window peak on Locus 1 (Figure S9) in the broader linkage block. However, these genes occurred well outside the *F*_ST_ window peak, and were spread among dozens of other genes that did not show habitat or temperature driven differences in gene expression.

### 4. Symbiont community composition with ITS2 metabarcoding

Despite the pronounced physiological and environmental differences between habtiats, our metabarcoding approach targeting *Symbiodiniaceae* ITS2 region of rDNA revealed colonies from both lagoon and slope habitats associated exclusively with *Cladocopium* spp (Figure 5). An average of 106,213 reads (+/− 5,990 se) per sample were clustered into 62 distinct ITS2 sequence variants (Table S10). All colonies were dominated by ITS2 type C40, with various background levels of other *Cladocopium* ITS2 types (Figure 3B, Table S7). Analyses in *symportal* indicated that these distinct ITS2 variants form two distinct ITS2 type profiles in the data (Figure 3B), one of which occurred at high frequency in one of the slope sites. This ITS2 type profile was driven by a background ITS2 variant 5231_C, which occurred at moderate frequencies (~7%) in the majority of S1 colonies but was mostly absent in colonies from other sites. This specificity to *Cladocopium* is a common and rather unique pattern across Western Australia^37,41,65,81^, in contrast to corals from other regions that show pronounced shifts in symbiont species composition across environmental gradients^62,64,82,83^.

**Figure 5.**
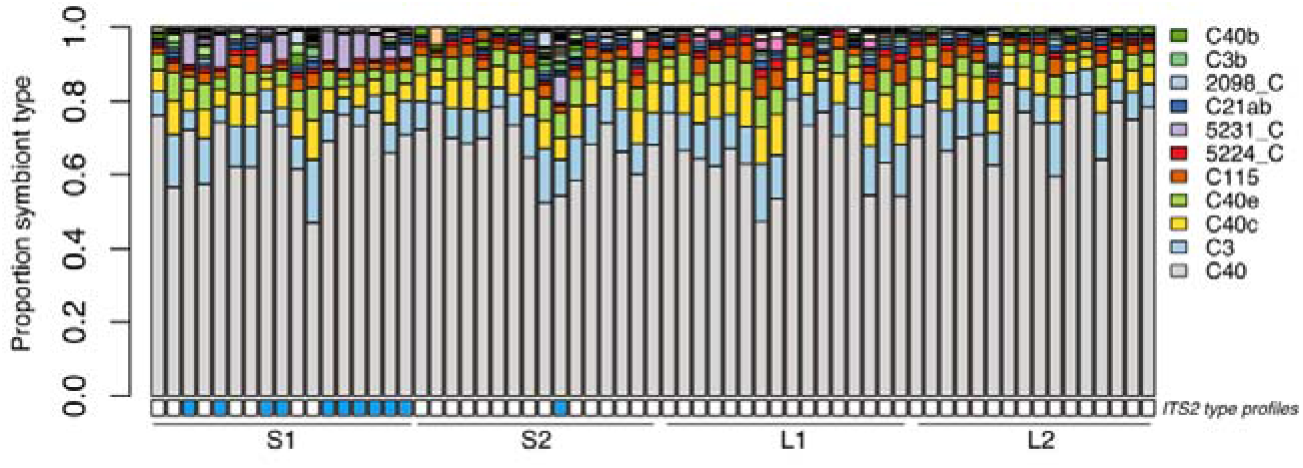
Symbiont community composition in A. tenuis from lagoon and slope habitats. Barplot of relative abundance of different symbiont ITS2 types found in corals from the two habitats at Clerke Reef based on ITS2 metabarcodiong. Each vertical bar represents an individual colony. Only the most abundant ITS2 variants are displayed in legend. Colored boxes below the plot show the distribution of the two symbiont ‘ITS2 type profiles’ identified in the data using symportal^66^.

### 5. Conclusions

Ecological divergence among populations of coral will be a key driver in their capacity to adapt to a rapidly changing environment. This is particularly true for isolated reef systems, which are largely reliant on local standing genetic variation to adapt to climate-related stressors. In Western Australia, the offshore oceanic atolls of the Rowley Shoals are isolated in space and time, characterized by strong genetic differentiation with neighbouring reef systems and a skew in the allele frequency spectrum indicative of a recent bottleneck. Across the Clerke Reef atoll, we identified high gene flow between slope and lagoon habitats, but identified a number of genomic islands of divergence that exhibited signs of restricted gene flow and strong linkage disequilibrium, suggesting that they may play an important role in adaptation to the marginal lagoon environment. Acute heat stress experiments showed lagoon corals to be more resistant to heat stress than slope corals, and accompanying RNA seq data showed that this was in part achieved through a habitat-specific physiological shift that involved hundreds of genes, a large fraction of which were part of a larger heat stress response. Identifying the genomic regions that confer resilience to different stressors, and understanding the physiological trade-offs associated with those shifts, will allow the development of targeted genetic assays to rapidly screen corals for the complex traits needed to effectively survive climate change. Such information will prove to be invaluable to managers focused on protecting pockets of corals more resistant to climate change and their application in active interventions and coral reef restoration programs^14,22,23,31^. This study contributes to the growing number of studies documenting the complex mechanisms that facilitate survival of corals across pronounced environmental gradients, and showcases the utility of combining different sequencing approaches to unravel complex climate-related traits and to identify mechanisms that will help slow coral reef degradation in the coming decades.

## Supporting information

ESM1

ESM2

## Acknowledgements

This work was conducted as part of the North West Shoals to Shore Program, which is proudly supported by Santos as part of the company’s commitment to better understand WA’s marine environment. The authors also acknowledge the support of ARC Linkage Project LP160101508 to explore coral resilience. JU was supported by the Woodside Coral Reef Research Fellowship.

