## Supplementary material for "Mechanisms of ecological divergence with gene flow in a reef-building coral on an isolated atoll in Western Australia": ESM1

**Electronic Supplementary Material 1**

Table S1 Two-week time-series temperature and current data from lagoon and slope habitat at Clerke Reef (.csv file).

Table S2 Microsatellite genotypes to confirm spring lineage (.csv file).

Table S3 Mapping metrics using BWA to *A. tenuis* assembly.

Table S4 List of genes with protein BLAST annotation for outlier Locus 1.

Table S5 List of genes with protein BLAST annotation for outlier Locus 3.

Table S6 Normalized gene expression counts matrix for heated and control samples (.csv).

Table S7 Results from differential gene expression analyses with DeSeq2 between colonies collected from lagoon and slope habitats after two days acclimation in a common garden (.csv).

Table S8 Results from differential gene expression analyses with DeSeq2 between heated and control treatments across all samples (.csv).

Table S9 List of genes differentially expressed between habitats in control treatments that also responded to acute heat stress.

Table S10 Counts matrix for the 62 ITS2 sequence variants recovered from colonies of A. tenuis from lagoon and slope habitats at Clerke Reef (.csv file).

Figure S1 Time-series temperature data (10-minute intervals) from lagoon and slope habitats at Clerke Reef collected across the 2017/2018 summer months. See ESM2 for fine-scale resolution across a two-week window in 2019.

Figure S2 Histogram of mean sequence coverage per base pair per sample for the entire dataset, and boxplot showing how those values varied by habitat and reef.

Figure S3 Initial screening of individuals using PCangsd. Dendrogram on left is based on a Euclidean distance matrix and shows samples to cluster strongly by reef system. The three outlier individuals that were removed from the dataset are indicated by red points. Two of these outliers shared the same MLG and were deemed clones. Plot on the right displays first two principal components and shows samples to cluster strongly by reef system.

Figure S4 TCS network based on extracted consensus sequences for each sample across the complete mitochondrion.

Figure S5 Minor allele frequencies in lagoon and slope samples from Rowley Shoals across Locus 1 and Locus 3.

Figure S6 Loading along the first principal component for Locus 1 and Locus 3 in colonies from Lagoon and Slope habitats

Figure S7 Visual scoring of colonies using the CoralWatch Coral Health Chart from the two habitats following acute heat stress.

Figure S8 Initial screening of genotypes from RNAseq experiment. Scatter plot of the first two principal components for all genotypes used in experimental heat stress tests (left) and for those after the autumn spawning lineage was removed (right). Colour indicates habitat or origin (red-lagoon, blue-slope).

Figure S9 Plot of F_ST_ for each SNP and in 100kb windows (red line) across Locus 1. Genic regions are highlighted in grey, and ones that were differentially expressed between heated and control colonies, or between habitats, are highlighted in red.

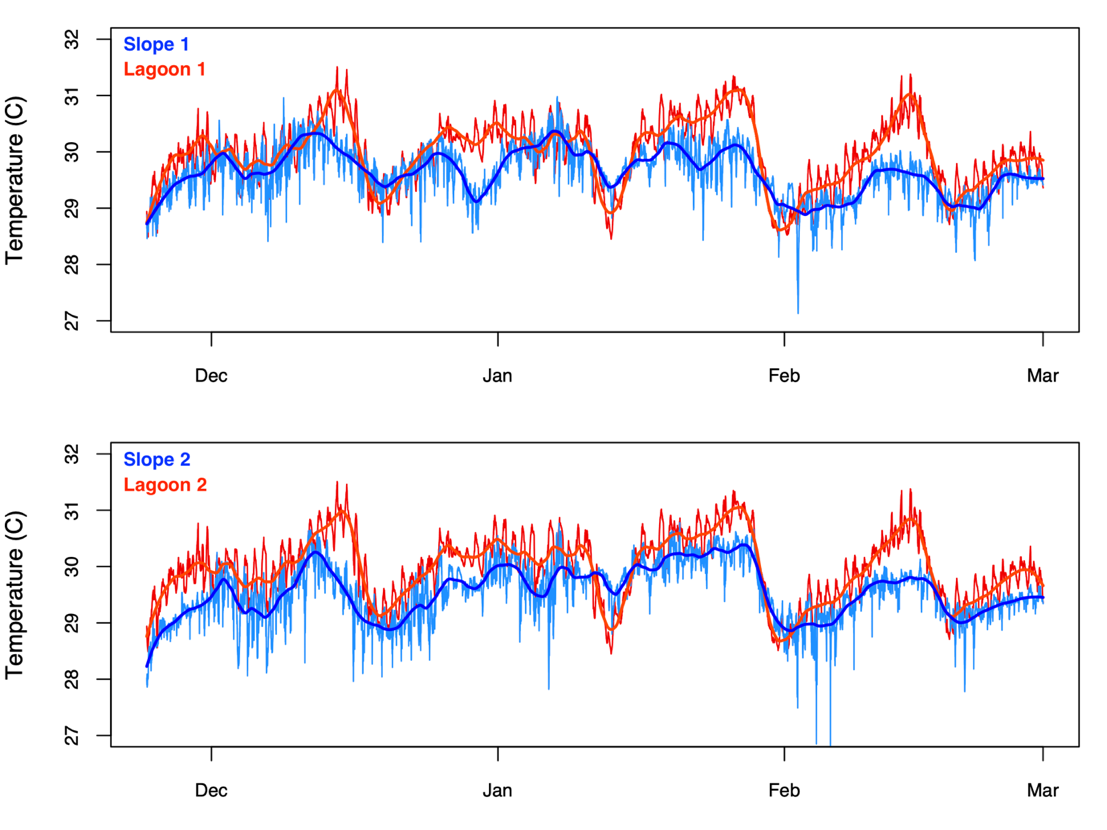

Figure S1 Time-series temperature data (10-minute intervals) from lagoon and slope habitats at Clerke Reef collected across the 2017/2018 summer month. See ESM2 for fine-scale resolution across a two-week window in 2019.
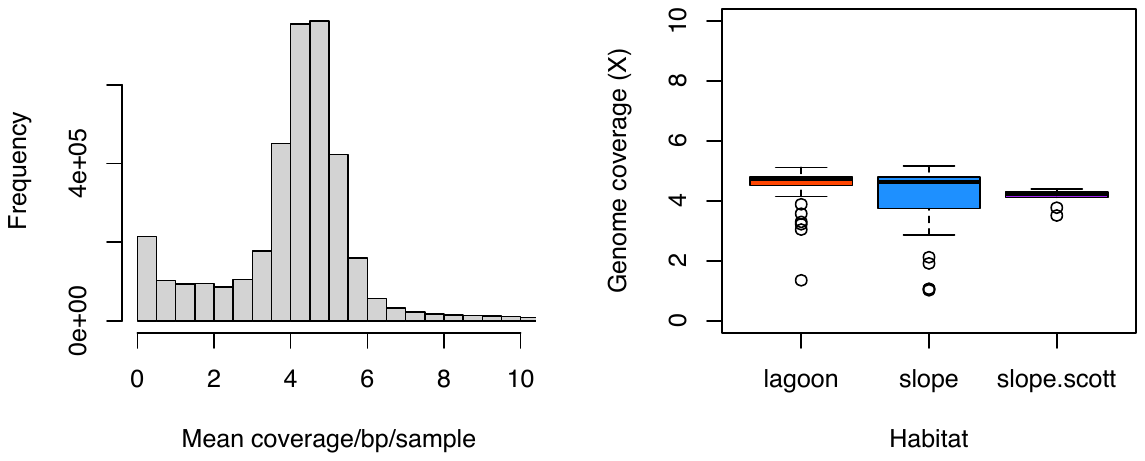

Figure S2 Histogram of mean sequence coverage per base pair per sample for the entire dataset, and boxplot showing how those values varied by habitat and reef.

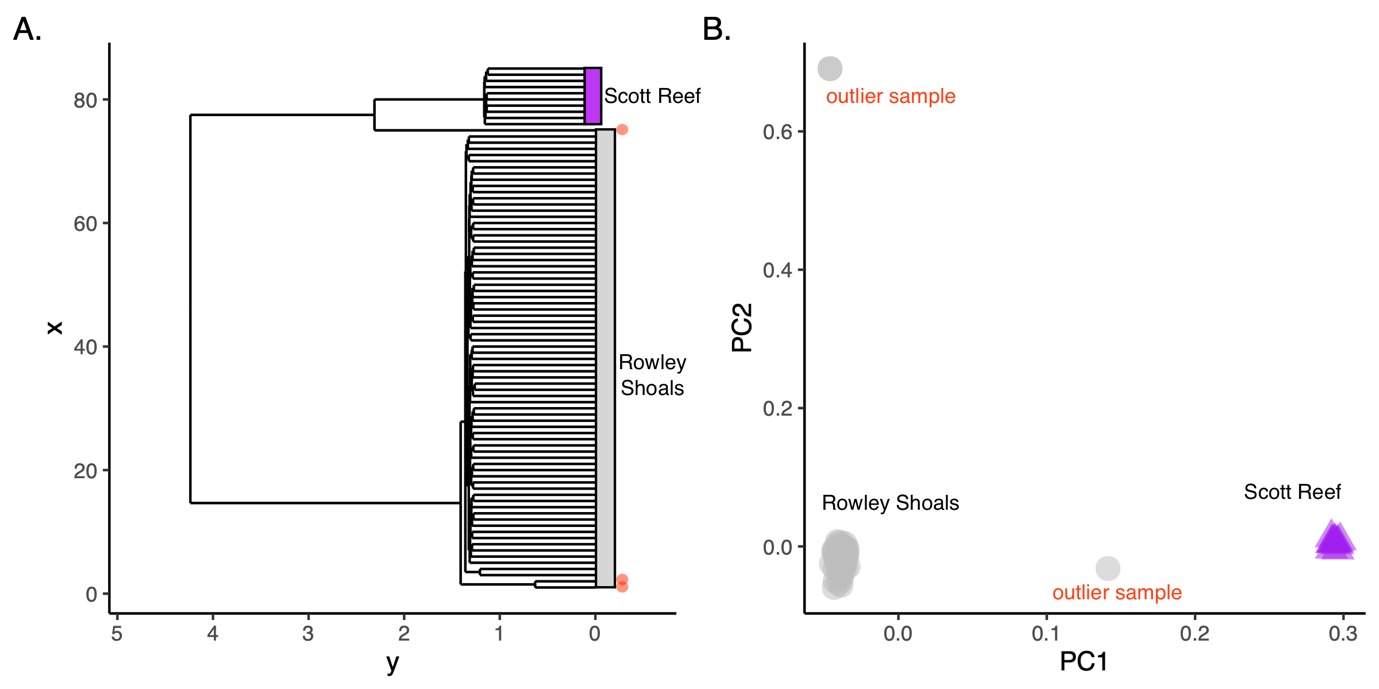

Figure S3 Initial screening of individuals using PCangsd. Dendrogram on left is based on a Euclidean distance matrix and shows samples to cluster strongly by reef system. The three outlier individuals that were removed from the dataset are indicated by red points. Two of these outliers shared the same MLG and were deemed clones. Plot on the right displays first two principal components component and shows samples to cluster strongly by reef system (Rowley Shoals and Scott Reef).

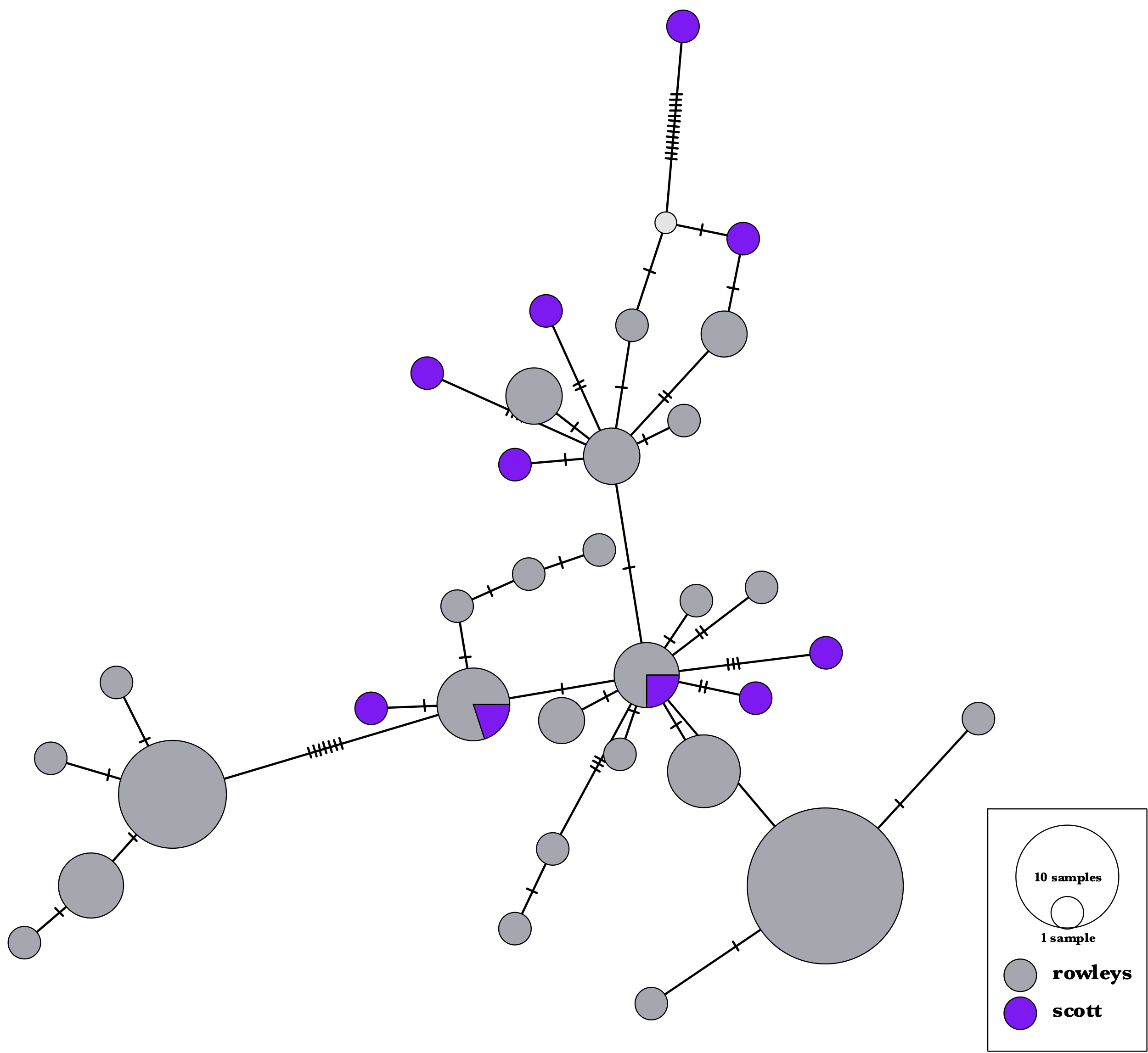

Figure S4 TCS network based on extracted consensus sequences for each sample across the complete mitochondrion

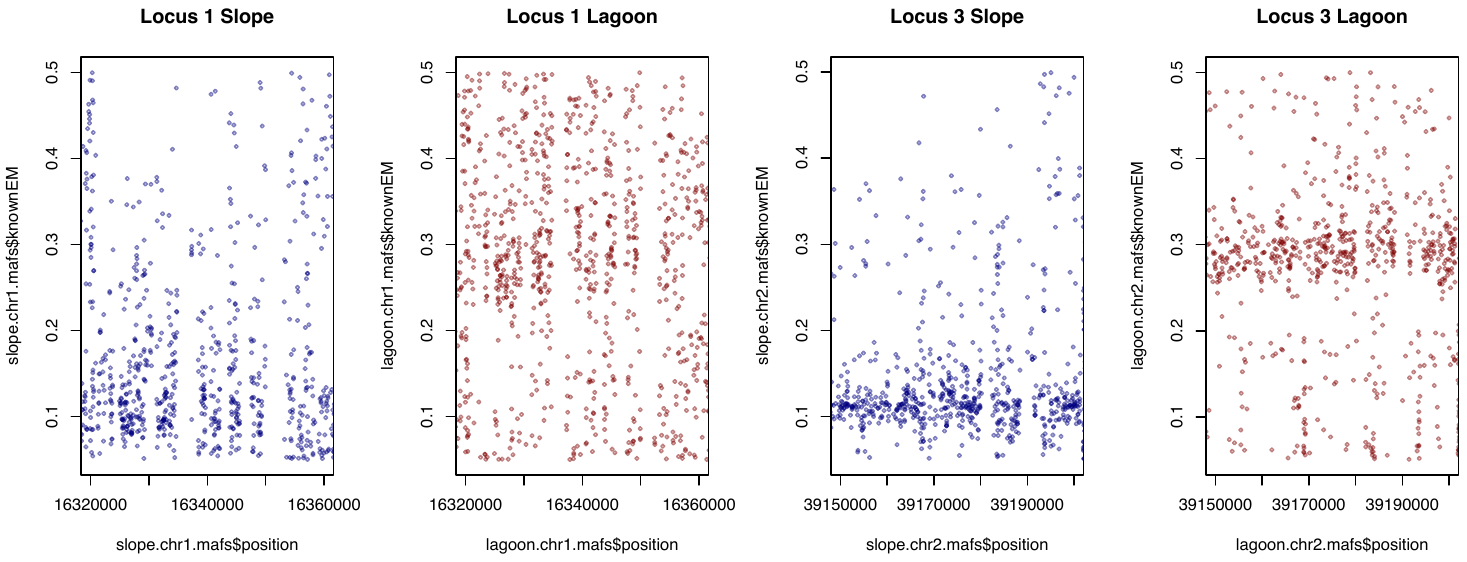

Figure S5 Minor allele frequencies in lagoon and slope samples from Rowley Shoals across Locus 1 and Locus 3.

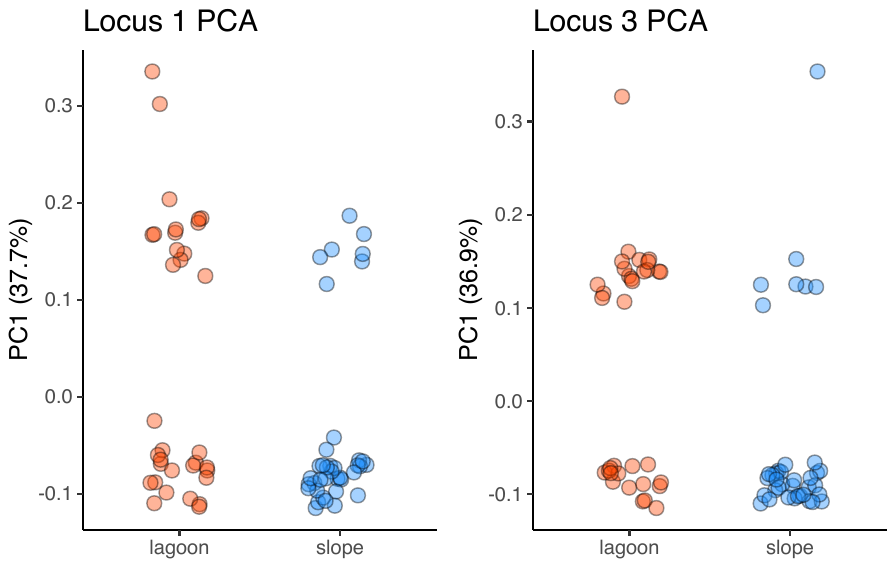

Figure S6 Loading along the first principal component for Locus 1 and Locus 3 in colonies from Lagoon and Slope habitats.

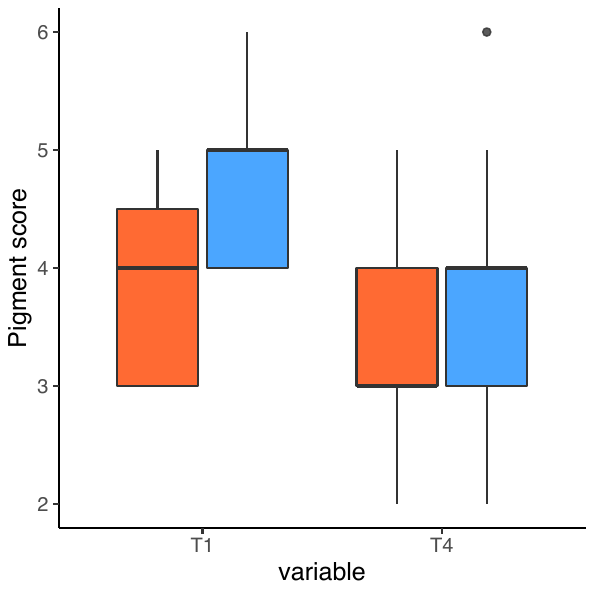

Figure S7 Visual scoring of colonies using the CoralWatch Coral Health Chart from the lagoon (red) and slope (blue) following acute heat stress.

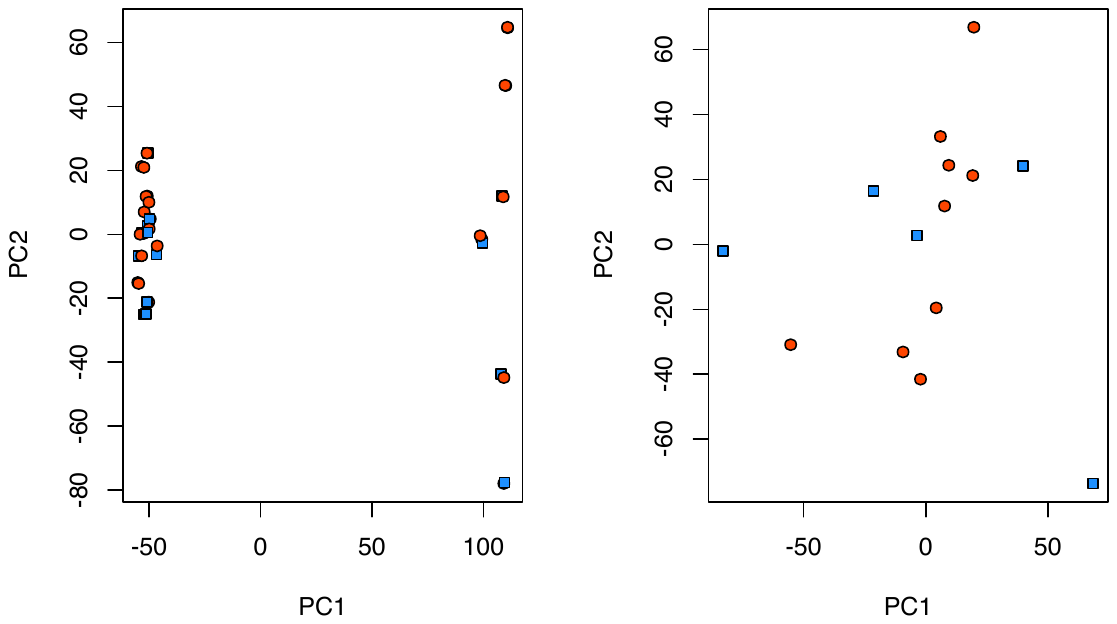

Figure S8 Initial screening of genotypes from RNAseq experiment. Scatter plot of the first two principal components for all genotypes used in experimental heat stress tests (left) and for those after the autumn spawning lineage was removed (right). Colour indicates habitat or origin (red-lagoon, blue-slope).

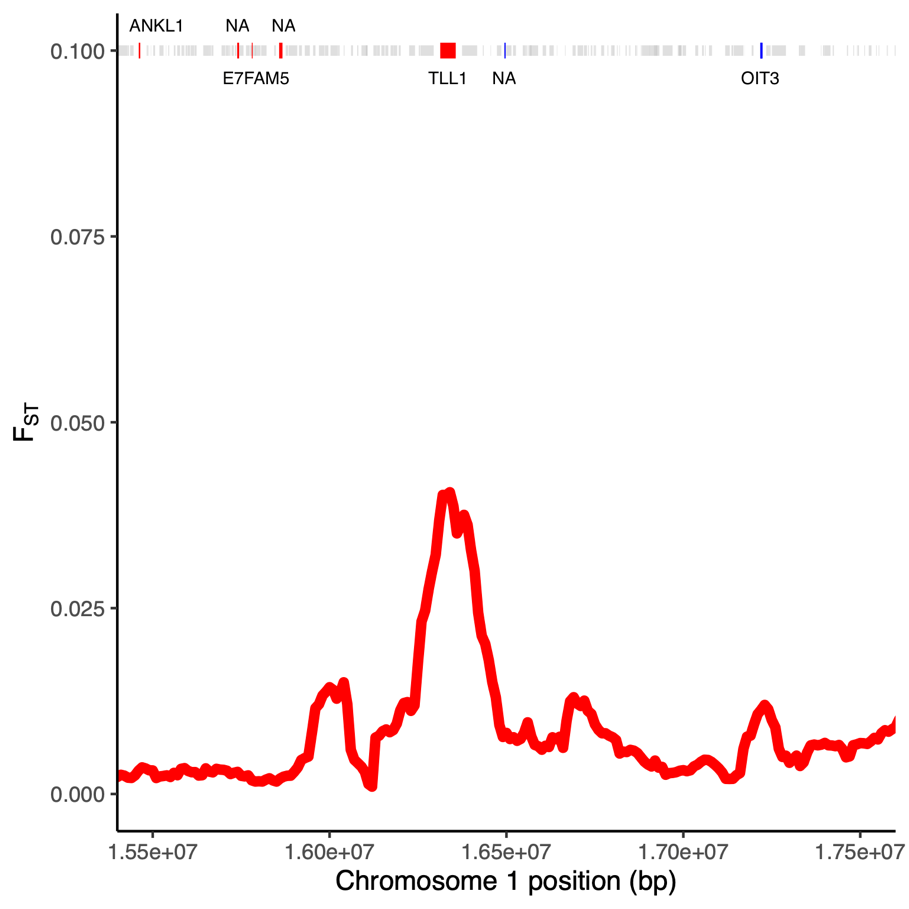

Figure S9 Plot of *F*_ST_ for each SNP and in 100kb windows (red line) across Locus 1. Genic regions are highlighted in grey, and ones that were differentially expressed between heated and control colonies are in red and between habitats in blue.
