## Supplementary material for "Mechanisms of ecological divergence with gene flow in a reef-building coral on an isolated atoll in Western Australia": ESM2

**Electronic Supplementary Material 2**

- **Temperature**

**Notes:** HOBOs were calibrated a week before the deployment. Response time for those instruments is of about 5 min and the accuracy is ±0.2°C accuracy.

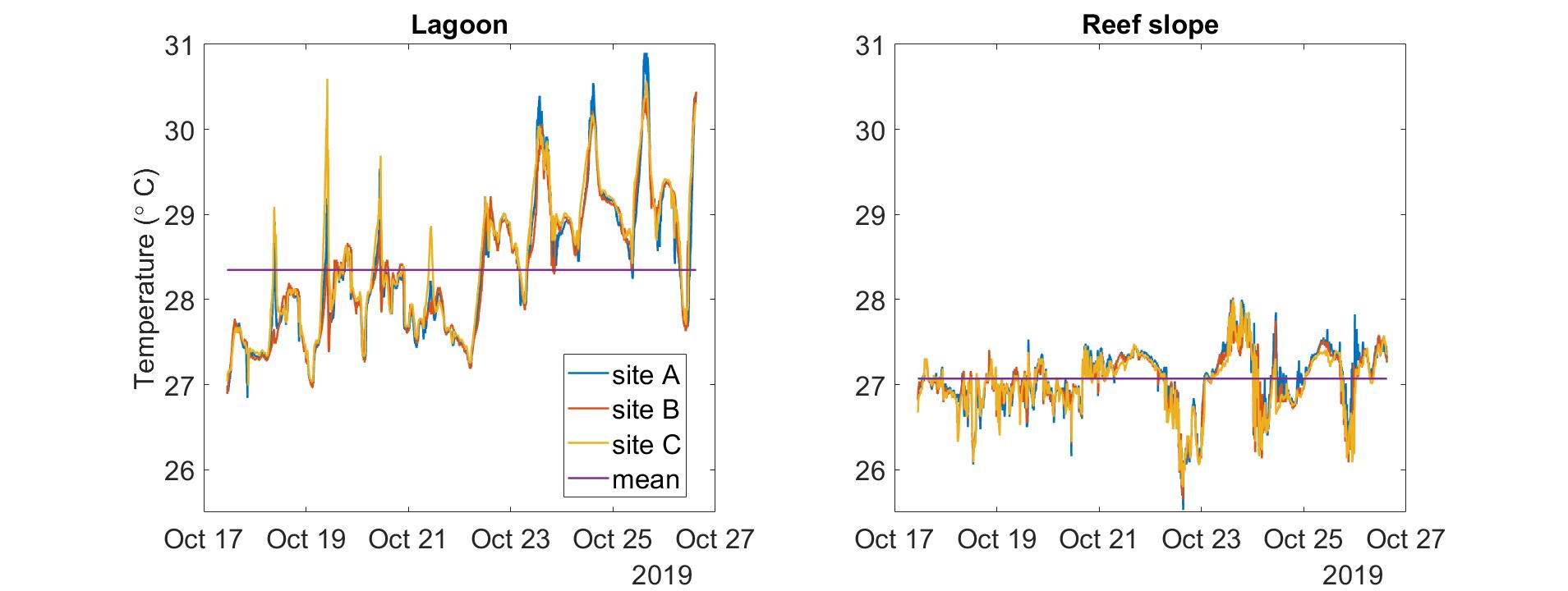

Figure 1: Lagoon and reef slope temperature at sites A, B and C at Clerke Reef

Lagoon:

| **Site** | **Mean(°C)** | **Variance(°C)** | **Mean daily variation(°C)** |
| --- | --- | --- | --- |
| Site A | 28.4 | 0.6 | 2 |
| Site B | 28.4 | 0.6 | 1.7 |
| Site C | 28.4 | 0.6 | 2 |

Reef slope

| **Site** | **Mean(°C)** | **Variance(°C)** | **Mean daily variation(°C)** |
| --- | --- | --- | --- |
| Site A | 27 | 0.1 | 1.1 |
| Site B | 27 | 0.1 | 1.1 |
| Site C | 27 | 0.1 | 1 |

- **Currents**

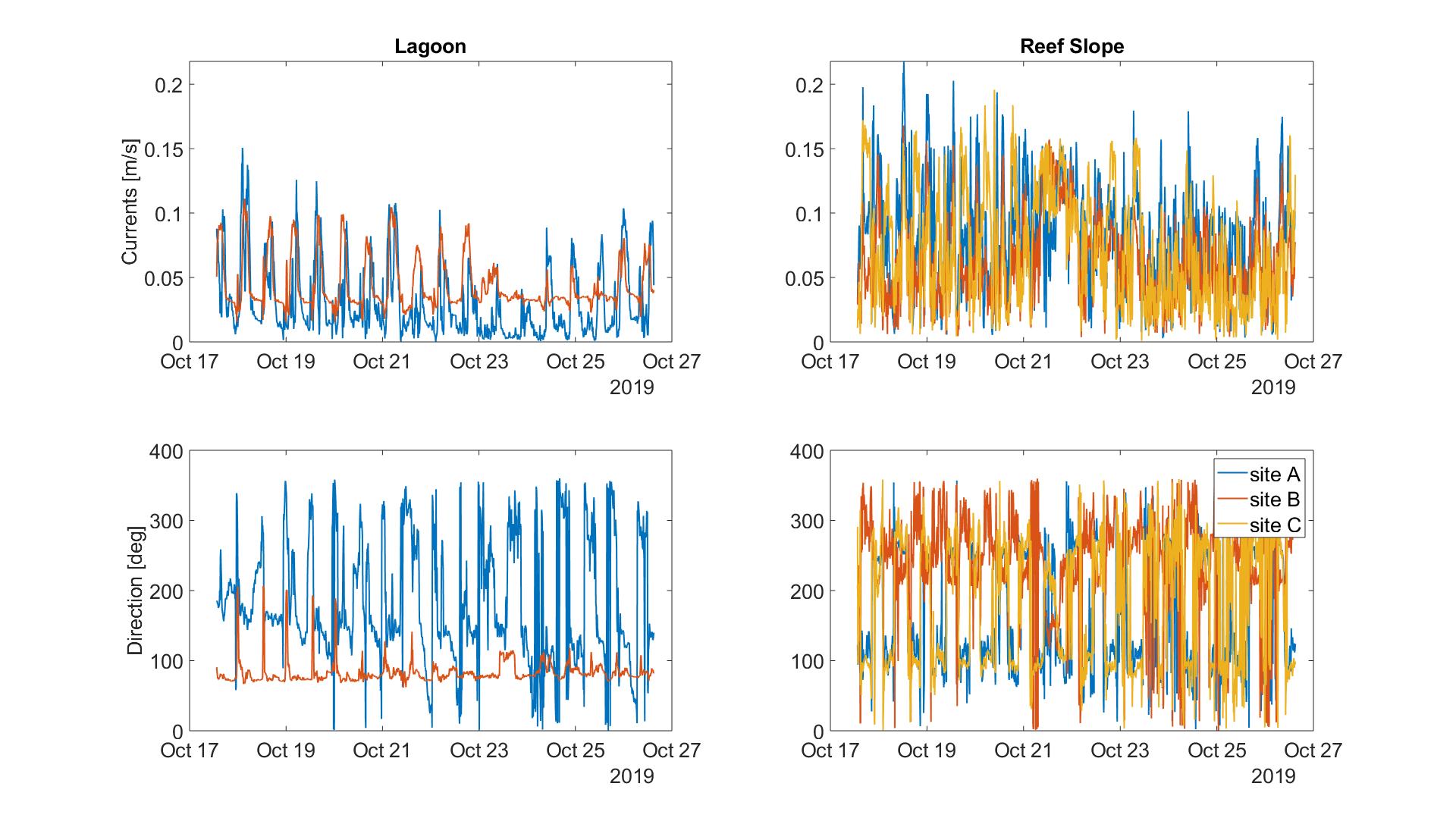

Lagoon:

| **Site** | **Mean [m/s]** | **Variance [m/s]** |
| --- | --- | --- |
| Site B | 0.03 | 7.10^-4^ |
| Site C | 0.04 | 3.10^-4^ |

Reef slope

| **Site** | **Mean [m/s]** | **Variance [m/s]** |
| --- | --- | --- |
| Site A | 0.08 | 0.002 |
| Site B | 0.06 | 0.001 |
| Site C | 0.06 | 0.002 |
